# Genomic and molecular characterisation of a *Klebsiella pneumoniae* clinical isolate resistant to meropenem-vaborbactam, imipenem-relebactam, and ceftazidime-avibactam

**DOI:** 10.1101/2025.08.11.669739

**Authors:** Yu Wan, Joshua L. C. Wong, Julia Sanchez-Garrido, Wen Wen Low, Jane F. Turton, Fabio Morecchiato, Ilaria Baccani, Kirsty Dodgson, Gian Maria Rossolini, Neil Woodford, Gad Frankel, Elita Jauneikaite, Danièle Meunier, Katie L. Hopkins

## Abstract

This article reports an unusual *Klebsiella pneumoniae* clinical isolate, KpMVR1, resistant to meropenem-vaborbactam, imipenem-relebactam, and ceftazidime-avibactam, and investigates the underlying genetic alterations using comparative genomics and molecular experiments.

Resistance to carbapenems and third-generation cephalosporins is increasing in *K. pneumoniae* globally, restricting therapeutic options. The β-lactam/β-lactamase inhibitor combinations are widely used to circumvent β-lactamase-mediated resistance. In 2021, isolate KpMVR1 was recovered from a hospitalised patient in England. Two additional isolates with the same variable-number tandem-repeat profile—KpMVS1, collected from the same patient 42 days before KpMVR1, and KpMVS2, from another patient in the same hospital—were susceptible to meropenem-vaborbactam, imipenem-relebactam, and ceftazidime-avibactam. Illumina and nanopore whole-genome sequencing and hybrid genome assembly were conducted for these three isolates. Annotated genome assemblies were compared to identify genetic variation, and mutagenesis experiments were performed to verify predicted functional alterations.

All isolates belonged to a novel clone ST8134 and carried *bla*_KPC-2_-like alleles (KpMVR1: *bla*_KPC-157_; KpMVS1 and KpMVS2: *bla*_KPC-2_) in presumptively conjugative plasmids. IS*Ec68* caused a frameshift mutation in KpMVR1’s *ompK36* gene, reducing the meropenem-vaborbactam and imipenem-relebactam susceptibility. KPC-157 demonstrated decreased hydrolysis of imipenem and ceftazidime when compared with KPC-2. KpMVR1 also encoded a disrupted transcriptional repressor MarR and a destabilising mutation in AcrB, a component of the AcrAB-TolC multidrug efflux pump. In conclusion, KpMVR1 harboured complex resistance-associated genetic alterations, with evidence for *in vivo* emergence of antimicrobial resistance. Our study underlines routine screening for resistant pathogens in vulnerable patients to guide antimicrobial chemotherapy as well as the need to characterise underlying resistance mechanisms to help assess the potential for onward transmission.

**Data summary:** Illumina and nanopore sequencing reads, hybrid genome assemblies, and anonymised metadata of isolates KpMVS1, KpMVR1, and KpMVS2 have been deposited in databases of the National Center for Biotechnology Information (www.ncbi.nlm.nih.gov) under BioProject accession PRJNA1084250, with BioSample accessions SAMN46778009 (KpMVS1), SAMN46778010 (KpMVR1), and SAMN46778011 (KpMVS2). The genome assemblies of these isolates have also been deposited in Pasteur Institute’s database for *K. pneumoniae* species complex (bigsdb.pasteur.fr/klebsiella/) under ids 75608 (KpMVS1), 75609 (KpMVR1), and 75610 (KpMVS2).

**Impact statement:** This is the first *bla*_KPC_-positive *K. pneumoniae* isolate referred to the UK’s national reference laboratory with resistance to three last-resort β-lactam/β-lactamase inhibitor combinations meropenem-vaborbactam, imipenem-relebactam, and ceftazidime-avibactam, implicating *in vivo* emergence of this unusual resistance profile during prolonged antimicrobial chemotherapy. This isolate belonged to a novel clone ST8134 and harboured a plasmid-borne *bla*_KPC-2_-like allele *bla*_KPC-157_. We identified complex genetic alterations in this isolate: chromosomal large deletions, point mutations, and an IS*Ec68*-induced loss-of-function truncation of the *ompK36* porin gene. We determined the impact of KPC-2, KPC-157, and the *ompK36* truncation on the susceptibility of *K. pneumoniae* to meropenem, meropenem-vaborbactam, imipenem, imipenem-relebactam, imipenem-avibactam, aztreonam, aztreonam-avibactam, ceftazidime, ceftazidime-avibactam, and cefiderocol. Our work underscores the need to monitor emerging resistance to beta-lactam/beta-lactamase inhibitor combinations in healthcare and to understand underlying resistance mechanisms for assessing the potential of resistance transmission.

## Introduction

Meropenem and imipenem are broad-spectrum carbapenem antimicrobials parenterally administered to treat serious bacterial infections, such as those caused by Enterobacterales producing extended-spectrum β-lactamases (ESBLs) or AmpC-type cephalosporinases (AmpCs) [1–3]. Ceftazidime, a third-generation parenteral cephalosporin, is also widely used for treating severe bacterial infections, although it can be hydrolysed by ESBLs and AmpCs [4, 5]. In Gram-negative bacteria, carbapenems and cephalosporins diffuse through outer-membrane porins and enter the periplasm, where they inactivate penicillin binding proteins, disrupting cell-wall synthesis with a bactericidal effect [6–8].

Combining β-lactams with β-lactamase inhibitors in antimicrobial chemotherapy is a widely employed strategy to circumvent β-lactamase-mediated resistance in bacteria. Vaborbactam, relebactam, and avibactam are non-β-lactam inhibitors of Ambler class A β-lactamases, such as ESBLs and *Klebsiella pneumoniae* carbapenemases (KPCs), as well as class C β-lactamases (AmpCs) [9]. Moreover, avibactam inhibits several class D β-lactamases such as OXA-48 and OXA-10 [10]. These inhibitors penetrate the outer membrane (OM) of Gram-negative bacteria via porins, blocking the active sites of β-lactamases in the periplasm [9, 11].

In the UK, meropenem-vaborbactam, imipenem-relebactam, and ceftazidime-avibactam are reserved for highly selected patients [12]. Resistance to one or more of these combination antimicrobials in clinical isolates of KPC-producing *Klebsiella pneumoniae* has been previously reported [13–15]. Underlying resistance mechanisms include overproduction of KPCs or the AcrAB-TolC multidrug efflux pump, as well as gain-of-function point mutations in the *bla*_KPC_ gene [9, 16–18]. In addition, the disruption or transcriptional downregulation of *ompK35* (*ompF*) and *ompK36* (*ompC*), which encode non-selective porins that facilitate the diffusion of β-lactams and β-lactamase inhibitors through the OM, has also been implicated [17–20].

Here, we report and characterise a clinically significant *K. pneumoniae* isolate exhibiting resistance to ceftazidime-avibactam, meropenem-vaborbactam, and imipenem-relebactam, analysed in the context of closely related isolates recovered in the same hospital. This was the first such isolate referred to the UK’s national reference laboratory for characterisation.

## Methods

### Isolate collection and phenotyping

The *K. pneumoniae* isolate KpMVS1 was recovered from a lung biopsy specimen of an inpatient (hereafter, Patient 1) admitted to an intensive care unit (ICU) in England in 2021. The second *K. pneumoniae* isolate, KpMVR1, was recovered from a groin wound of the same patient 42 days later. During this ICU stay, the patient received a broad range of antimicrobials, including meropenem-vaborbactam, ciprofloxacin, and gentamicin; however, ceftazidime-avibactam and imipenem-relebactam were not used.

Species identification of the isolates and carbapenemase gene screening were performed using matrix-assisted laser desorption/ionisation-time of flight (MALDI-ToF) method and the GeneXpert system (Cepheid, USA), respectively. Initial antimicrobial susceptibility testing (AST) was conducted by the hospital, with results interpreted according to the European Committee on Antimicrobial Susceptibility Testing (EUCAST) guidelines. Both isolates were referred to the Antimicrobial Resistance and Healthcare Associated Infections (AMRHAI) Reference Unit of the UK Health Security Agency (UKHSA) for variable number tandem repeat (VNTR) typing [21] and investigation of unusual antimicrobial resistance (AMR). Furthermore, a *bla*_KPC_-positive *K. pneumoniae* isolate KpMVS2 — recovered from a rectal swab of another patient (Patient 2) in the same hospital in 2020 during an outbreak investigation and sharing the same VNTR profile as KpMVS1 and KpMVR1 — was retrieved from AMRHAI’s culture collection for comparison. These three isolates were subjected to whole-genome sequencing (WGS) and AST for which minimum inhibitory concentrations (MICs) of 19 antimicrobials (Table 1) and diameters of cefiderocol inhibition zones were interpreted as per EUCAST clinical breakpoints v15.0 [22].

**Table 1.** Antimicrobial minimum inhibitory concentrations (MICs; mg/L) determined by UKHSA’s AMRHAI Reference Unit and susceptibility interpretations (as per EUCAST clinical breakpoints v15.0) of three *K. pneumoniae* clinical isolates. Abbreviations: MEM, meropenem; VAB, vaborbactam; IPM, imipenem; ETP, ertapenem; CET, ceftolozane; TZB, tazobactam; CFD, cefiderocol; FEP, cefepime; CAZ, ceftazidime; AVI, avibactam; CTX, cefotaxime; FOX, cefoxitin; TMC, temocillin; AMP, ampicillin; AMX, amoxicillin; CAV, clavulanate; PIP, piperacillin; ATM, aztreonam; AMK, amikacin; GEN, gentamicin; CST, colistin; CIP, ciprofloxacin; MB, monobactam. Susceptibility interpretations: R, resistant; I, susceptible, increased exposure; S, susceptible.

|  | Carbapenem |  |  |  | Cephalosporin |  |  |  |  |  |  | Penicillin |  |  |  | MB | Aminoglycoside |  | Others |  |
| --- | --- | --- | --- | --- | --- | --- | --- | --- | --- | --- | --- | --- | --- | --- | --- | --- | --- | --- | --- | --- |
| Isolate | MEM-VAB | MEM | IPM | ETP | CET-TZB | CFD* | FEP | CAZ | CAZ-AVI | CTX | FOX‡ | TMC‡ | AMP | AMX-CAV | PIP-TZB | ATM | AMK‡ | GEN‡ | CST‡ | CIP |
| KpMVR1 | >256 (R) | >16 (R) | >128 (R) | >4 (R) | 16 (R) | (S) | 4 (I) | 256 (R) | 16 (R) | 8 (R) | >64 | >128 | >32 (R) | >32 (R) | >64 (R) | 16 (R) | 2 | 0.5 | ≤0.5 | >4 (R) |
| KpMVS1 | 0.064 (S) | >16 (R) | 64 (R) | >4 (R) | >16 (R) | (S) | >32 (R) | 128 (R) | 1 (S) | 64 (R) | >64 | 32 | >32 (R) | >32 (R) | >64 (R) | >32 (R) | ≤1 | ≤0.25 | ≤0.5 | ≤0.125 (S) |
| KpMVS2 | 0.032 (S) | 16 (R) | 16 (R) | >4 (R) | >16 (R) | (R) | >32 (R) | 64 (R) | 1 (S) | 16 (R) | 32 | 8 | >32 (R) | >32 (R) | >64 (R) | >32 (R) | ≤1 | ≤0.25 | 1 | ≤0.125 (S) |

\* CFD susceptibility was determined using disc diffusion. ‡ Interpretations were not available according to EUCAST guidelines.

### Whole-genome sequencing

Genomic DNA of each isolate was extracted from an overnight culture using the GeneJET Kit (ThermoFisher Scientific, UK) as per the manufacturer’s protocol. Short-read sequencing was conducted on a HiSeq 2500 system (Illumina, USA) by UKHSA’s Colindale Sequencing Laboratory following its paired-end 101-bp protocol. Long-read sequencing was performed on MinION R9.4.1 flow cells (Oxford Nanopore Technologies [ONT], UK), with libraries prepared using the ONT Rapid Barcoding Kit SQK-RBK004.

### Bioinformatics analysis

Illumina reads were trimmed and filtered with Trimmomatic v0.39 for a minimum per-read quality of Phred Q30 and minimum length of 50 bp [23]. Fast-mode basecalling and de-multiplexing of nanopore reads was conducted by guppy v4 (ONT). Nanopore reads were then trimmed and filtered for a minimum per-read quality of Q10 and minimum length of 1 kbp using fastp v0.23.4 [24]. For species confirmation and contamination assessment, taxonomical profiling of processed Illumina and nanopore reads were performed using Kraken v2.1.3, bracken v2.8, and a standard Kraken database built in September 2023 [25, 26].

Genomes of KpMVR1 and KpMVS2, with estimated nanopore read depths of 185× and 243×, respectively, were assembled using hybracter v0.5.0 (assemblers: Flye v2.9.3 and plassembler v1.5.0; sequence re-orientator: dnaapler v0.5.1; long-read polisher: medaka v1.8.0, short-read polishers: pypolca v0.2.1 and polypolish v0.5.0) [27–32]. For KpMVS1, which had nanopore reads with an estimated depth of 64×, the chromosome and plasmid sequences were assembled using Raven v1.8.3 and plassembler, respectively, and polished with nanopore and Illumina reads as for KpMVR1 and KpMVS2. Evaluation of contamination and completeness of genome assemblies were conducted with CheckM2 v1.0.2 and its database Uniref100/KO [33]. The average fold-coverage of each contig was estimated from Illumina and nanopore reads, respectively, using mosdepth v0.3.9 [34].

The genome assemblies were annotated using bakta v1.9.2 and its standard database v5.1 [35]. Multi-locus sequence typing, serotype prediction, and virulence-factor detection were performed using Kleborate v3.1.3, which incorporated Kaptive v3.1.0 [36, 37]. AMR genes were detected using AMRFinderPlus v3.12.8 with a minimum query coverage of 80% [38]. Clustered regularly interspaced short palindromic repeats (CRISPR) and CRISPR-associated (Cas) genes in chromosomes were predicted using CRISPRCasFinder [39]. For plasmids, replicon types were determined at a minimum of 80% nucleotide identity and coverage using PlasmidFinder v2.1 [40] and the mobility was predicted using mob_typer of MOB-suite v3.1.8 [41]. The fold-coverage of each KPC-encoding plasmid was divided by that of its host’s chromosome to estimate the plasmid copy number. Transposons and insertion sequences (ISs) were identified using TnCentral Blast (blastn) and ISFinder, respectively [42, 43].

Chromosome and plasmids of KpMVR1 and KpMVS2 were compared against those of KpMVS1 using minimap v2.26 [44]. Identified genetic variants were annotated using snpEff v5.2 [45]. Gene Ontology terms were predicted from amino acid sequences using InterProScan v5.69-101.0 [46] with sequence alignments filtered for ≥60% query coverages. Impacts of point mutations on protein stability were predicted from wild-type protein structures in the UniProt database using Missense3D and DDMut [47–49]. The three-dimensional structure of the plasmid-encoded donor OM protein TraN was compared between KPC-encoding, IncFII-carrying plasmids pKpMVS1_1, pKpMVR1_1, and pKpMVS2_1 following the approach developed by Low *et al* (Supplementary methods) to estimate the impact of TraN alterations on the conjugation specificity and efficiency [50]. Comparison and annotations of these three plasmids were visualised using BRIG v0.95 and Proksee [51, 52]. Gene synteny was illustrated using R package gggenes [53]. Genetic alterations found in both KpMVR1 and KpMVS2 were considered unlikely to confer the unique AMR profiles of KpMVR1 and were therefore excluded from further investigation.

### Functional assessment

To experimentally determine and compare the impacts of *bla*_KPC-2_ and *bla*_KPC-157_ on β-lactam susceptibility in *K. pneumoniae*, KpMVS1’s KPC-2-encoding plasmid pKpMVS1_1, was introduced into the plasmid-free *K. pneumoniae* laboratory strain ICC8001 (MICs: meropenem, ≤0.06 mg/L; imipenem, 0.25 mg/L; aztreonam, ≤0.125 mg/L; ceftazidime and ceftazidime-avibactam, 0.25 mg/L) through conjugation, resulting in a transconjugant ICC8001_KPC-2_ [54]. Transgenic isolates ICC8001_KPC-157_ and KpMVS1_KPC-157_ were derived from ICC8001_KPC-2_ and KpMVS1, respectively, by substituting the *bla*_KPC-2_ allele with *bla*_KPC-157_. Moreover, isolates ICC8001KPC-2/ΔompK36 and ICC8001KPC-157/ΔompK36 were derived from ICC8001_KPC-2_ and ICC8001_KPC-157_, respectively, through seamless, markerless homologous recombination using mutagenesis vectors and a lambda-red based recombination system generated in previous work [54].

To predict the presence/absence of OmpK36 in the OM of KpMVR1, Sec-dependent signal peptides and their cleavage site in translated *ompK36* alleles were compared between KpMVS1 and KpMVR1 using SignalP v6.0 [55]. To validate the prediction, purification of OM proteins was performed by resuspending an overnight LB-Miller culture (VWR, USA) of each isolate in 1M HEPES (pH 7.4) and sonicating at 25% amplitude for 10 bursts of 10 seconds on, 15 seconds off each (Model 705 Sonic Dismembrator, Fisher Scientific). Isolates ICC8001 and its *ompK36*-knockout derivative, ICC8001_ΔompK36_, were included as positive and negative controls, respectively. After separating cellular debris by centrifugation, OM proteins were obtained by centrifugation at 14,000×g for 30 mins and resuspended in 2% sarcosine/HEPES for 30 mins at room temperature. All steps were performed at 4°C on ice to preserve protein integrity unless otherwise indicated. For visualisation, 10 μg protein per isolate was separated by sodium dodecyl sulfate–polyacrylamide gel electrophoresis (SDS-PAGE) using 12% acrylamide gels and was stained with Coomassie solution (Sigma-Aldrich, USA) and imaged on a ChemiDoc XRS+ (Bio-Rad, USA).

Both progenitor isolates (KpMVS1 and KpMVR1) and the five transgenic isolates (KpMVS1_KPC-157_, ICC8001KPC-2, ICC8001KPC-157, ICC8001KPC-2/ΔompK36, and ICC8001KPC-157/ΔompK36) were tested for susceptibility to meropenem, meropenem-vaborbactam, imipenem, imipenem-relebactam, imipenem-avibactam, ceftazidime, ceftazidime-avibactam, aztreonam, aztreonam-avibactam, and cefiderocol (in iron-depleted medium) by the reference broth microdilution method as per the EUCAST guidance [56, 57]. Any MIC change above a two-fold difference between two isolates was considered notable.

## Results

### Phenotypes of isolates

Isolates KpMVS1 and KpMVR1, obtained 42 days apart from Patient 1 with recurrent *K. pneumoniae* infections, and KpMVS2, obtained from Patient 2 in the same hospital, were identified as *K. pneumoniae* by both MALDI-ToF and WGS. In the ICU where KpMVS1 and KpMVR1 were recovered, all patients were screened for carriage of carbapenemase-producing Enterobacterales (CPE) on admission using PCR, and the resistance profile of KpMVR1 was unique among all identified CPE isolates from the ICU during Patient 1’s stay.

Based on AST results from the AMRHAI Reference Unit (Table 1), KpMVR1 was resistant to meropenem-vaborbactam (MIC>256 mg/L), ceftazidime-avibactam (MIC=16 mg/L), and ciprofloxacin (MIC>4 mg/L), whereas KpMVS1 and KpMVS2 were susceptible to these antimicrobial agents (MICs: meropenem-vaborbactam ≤0.064 mg/L; ceftazidime-avibactam 1 mg/L; ciprofloxacin ≤0.125 mg/L). Notably, KpMVS2 was resistant to cefiderocol. Further AST discovered that the imipenem-relebactam MIC of KpMVR1 (512 mg/L, resistant) was 2048 times that of KpMVS1 (0.25 mg/L, susceptible) (Table 2). Moreover, KpMVR1 exhibited a >4-fold increase in the temocillin MIC (>128 mg/L) and a >8-fold reduction in the cefepime MIC (4 mg/L, susceptible, increased exposure) compared with KpMVS1 (temocillin: 32 mg/L; cefepime: >32 mg/L, resistant).

**Table 2.**
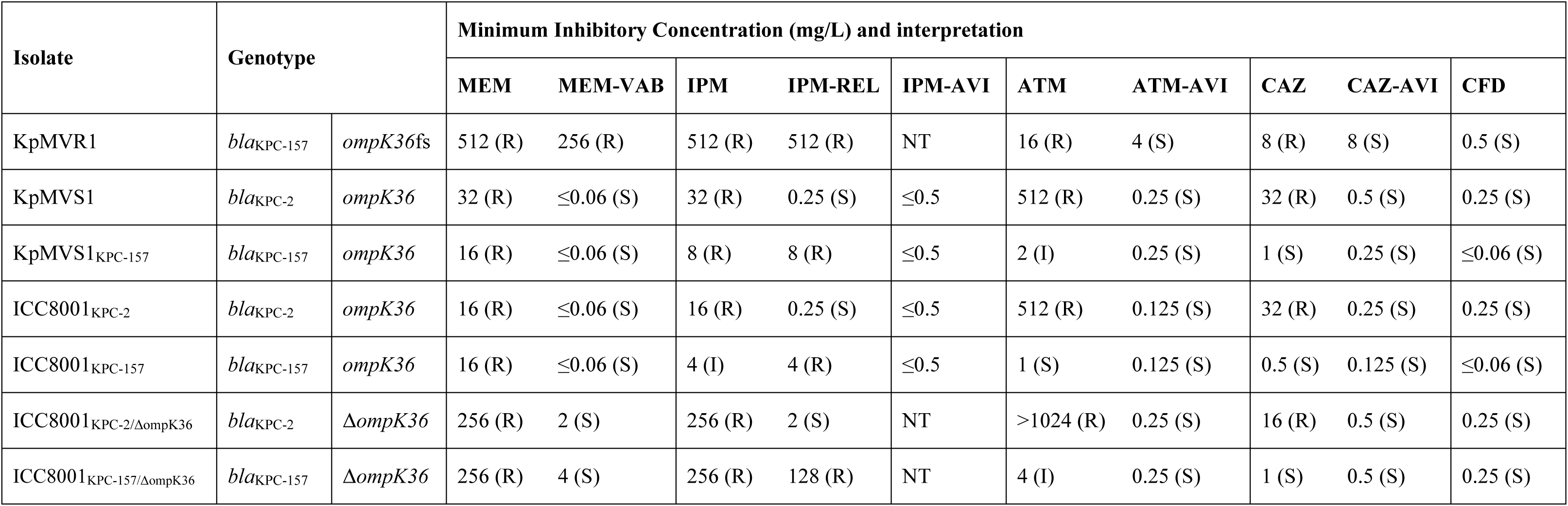
Minimum inhibitory concentrations of beta-lactam antimicrobials and susceptibility interpretations (as per EUCAST clinical breakpoints v15.0, where applicable) of progenitor and transgenic *K. pneumoniae* isolates determined in the experiments for functional assessment. Subscripts in isolate names indicate transgenic isolates and corresponding genotypes. Abbreviations: MEM, meropenem; VAB, vaborbactam; IPM, imipenem; REL, relebactam; ATM, aztreonam; AVI, avibactam; CAZ, ceftazidime; CFD, cefiderocol, tested in iron-depleted Mueller Hinton broth; NT, not tested. Interpretations of antimicrobial susceptibility: R, resistant; I, susceptible upon increased antimicrobial exposure; S, susceptible. Notations: *ompK36*fs, frameshifted *ompK36*; Δ*ompK36*, deletion of *ompK36*.

| Isolate | Genotype |  | Minimum Inhibitory Concentration (mg/L) and interpretation |  |  |  |  |  |  |  |  |  |
| --- | --- | --- | --- | --- | --- | --- | --- | --- | --- | --- | --- | --- |
|  |  |  | MEM | MEM-VAB | IPM | IPM-REL | IPM-AVI | ATM | ATM-AVI | CAZ | CAZ-AVI | CFD |
| KpMVR1 | *bla*<sub>KPC-157</sub> | *ompK36fs* | 512 (R) | 256 (R) | 512 (R) | 512 (R) | NT | 16 (R) | 4 (S) | 8 (R) | 8 (S) | 0.5 (S) |
| KpMVS1 | *bla*<sub>KPC-2</sub> | *ompK36* | 32 (R) | ≤0.06 (S) | 32 (R) | 0.25 (S) | ≤0.5 | 512 (R) | 0.25 (S) | 32 (R) | 0.5 (S) | 0.25 (S) |
| KpMVS1<sub>KPC-157</sub> | *bla*<sub>KPC-157</sub> | *ompK36* | 16 (R) | ≤0.06 (S) | 8 (R) | 8 (R) | ≤0.5 | 2 (I) | 0.25 (S) | 1 (S) | 0.25 (S) | ≤0.06 (S) |
| ICC8001<sub>KPC-2</sub> | *bla*<sub>KPC-2</sub> | *ompK36* | 16 (R) | ≤0.06 (S) | 16 (R) | 0.25 (S) | ≤0.5 | 512 (R) | 0.125 (S) | 32 (R) | 0.25 (S) | 0.25 (S) |
| ICC8001<sub>KPC-157</sub> | *bla*<sub>KPC-157</sub> | *ompK36* | 16 (R) | ≤0.06 (S) | 4 (I) | 4 (R) | ≤0.5 | 1 (S) | 0.125 (S) | 0.5 (S) | 0.125 (S) | ≤0.06 (S) |
| ICC8001<sub>KPC-2/ΔompK36</sub> | *bla*<sub>KPC-2</sub> | Δ*ompK36* | 256 (R) | 2 (S) | 256 (R) | 2 (S) | NT | >1024 (R) | 0.25 (S) | 16 (R) | 0.5 (S) | 0.25 (S) |
| ICC8001<sub>KPC-157/ΔompK36</sub> | *bla*<sub>KPC-157</sub> | Δ*ompK36* | 256 (R) | 4 (S) | 256 (R) | 128 (R) | NT | 4 (I) | 0.25 (S) | 1 (S) | 0.5 (S) | 0.25 (S) |

### Genetic characteristics of isolates

All three isolates belonged to *K. pneumoniae* clone ST8134, a novel single-locus variant of ST240, and were predicted to share the O1αβ,2β O-antigen type and K62 capsular polysaccharide type. Sequence lengths, plasmid replicons, and AMR genes determined in hybrid genome assemblies are summarised in Table 3. KpMVR1 and KpMVS1 shared the same plasmid types IncFII/repB(R1701), IncFII(pMET), Col(pHAD28), and Col(pHAD28)/Col440II, whereas KpMVS2 possessed unique plasmid types IncFII/IncR and IncFII(pKP91)/FIB(K).

**Table 3.** Genetic characteristics of the three *K. pneumoniae* clinical isolates. Abbreviation: AMR, antimicrobial resistance. Each hit of plasmid replicons covered the full length of its reference sequence in the PlasmidFinder database.

| Isolate | Sequence | Category | Length (bp) | Plasmid type | Nucleotide identity to template | Plasmid mobility | AMR gene |
| --- | --- | --- | --- | --- | --- | --- | --- |
| KpMVS1 | KpMVS1 | Chromosome | 5,404,514 |  |  |  | *bla<sub>SHV-36</sub>, fosA10* |
|  | pKpMVS1_1 | Plasmid | 111,397 | IncFII/repB(R1701) | IncFII(pKP91): 100%; repB(R1701): 99.52% | Conjugative | *bla<sub>KPC-2</sub>* |
|  | pKpMVS1_2 | Plasmid | 41,868 | IncFII(pMET) | IncFII(pMET): 98.09% | Non-mobilisable |  |
|  | pKpMVS1_3 | Plasmid | 4,809 | Col(pHAD28) | Col(pHAD28): 92.37% | Non-mobilisable |  |
|  | pKpMVS1_4 | Plasmid | 4,439 | Col(pHAD28)/Col440II | Col(pHAD28): 93.13%; Col440II: 97.52% | Mobilisable |  |
|  | pKpMVS1_5 | Plasmid | 3,258 | Unknown | Not detected | Non-mobilisable |  |
|  | pKpMVS1_6 | Plasmid | 1,917 | Col(pHAD28) | Col(pHAD28): 100% | Mobilisable |  |
| KpMVR1 | KpMVR1 | Chromosome | 5,292,801 |  |  |  | *bla<sub>SHV-36</sub>, fosA10* |
|  | pKpMVR1_1 | Plasmid | 111,174 | IncFII/repB(R1701) | IncFII(pKP91): 100%, repB(R1701): 99.52% | Conjugative | *bla<sub>KPC-157</sub>* |
|  | pKpMVR1_2 | Plasmid | 41,868 | IncFII(pMET) | IncFII(pMET): 98.09% | Non-mobilisable |  |
|  | pKpMVR1_3 | Plasmid | 4,809 | Col(pHAD28) | Col(pHAD28): 92.37% | Non-mobilisable |  |
|  | pKpMVR1_4 | Plasmid | 4,439 | Col(pHAD28)/Col440II | Col(pHAD28): 93.13%; Col440II: 97.52% | Mobilisable |  |
|  | pKpMVR1_5 | Plasmid | 3,258 | Unknown | Not detected | Non-mobilisable |  |
|  | pKpMVR1_6 | Plasmid | 1,917 | Col(pHAD28) | Col (pHAD28): 100% | Mobilisable |  |
| KpMVS2 | KpMVS2 | Chromosome | 5,354,507 |  |  |  | *bla<sub>SHV-36</sub>, fosA10* |
|  | pKpMVS2_1 | Plasmid | 116,795 | IncFII/IncR | IncFII(pKP91): 100%; IncR: 99.6% | Conjugative | *bla<sub>KPC-2</sub>* |
|  | pKpMVS2_2 | Plasmid | 41,868 | IncFII(pMET) | IncFII(pMET): 98.09% | Non-mobilisable |  |
|  | pKpMVS2_3 | Plasmid | 4,808 | Col(pHAD28) | Col (pHAD28): 92.37%* | Non-mobilisable |  |
|  | pKpMVS2_4 | Plasmid | 4,187 | Col(pHAD28) | Col (pHAD28): 92.37%* | Non-mobilisable |  |
|  | pKpMVS2_5 | Plasmid | 3,258 | Unknown | Not detected | Non-mobilisable |  |
|  | pKpMVS2_6 | Plasmid | 1,917 | Col(pHAD28) | Col (pHAD28): 100% | Mobilisable |  |
|  | pKpMVS2_7 | Plasmid | 240,297 | IncFII(pKP91)/IncFIB(K) | IncFII(pKP91): 84.98%; IncFIB(K): 98.93% | Conjugative |  |

\* Two hits of the Col(pHAD28) template sequence in the PlasmidFinder database differed between pKpMVS2\_3 and pKpMVS2\_4 by seven nucleotide substitutions (95% nucleotide identity) despite their same percent identity to the template.

KpMVS1 carried *bla*_KPC-2_ on the 111.4 kbp IncFII(pKP91)/repB(R1701) plasmid pKpMVS1_1. A plasmid of the same type was identified in KpMVR1 (pKpMVR1_1, 111.2 kbp) and carried *bla*_KPC-157_, which differed from *bla*_KPC-2_ by a single missense mutation (392A>G) resulting in an N131S amino acid substitution within the enzyme’s active site [58], where N131 bounds to relebactam, avibactam, and vaborbactam through a hydrogen bond [59–61]. Notably, pKpMVR1_1 differed from pKpMVS1_1 by 285 nucleotide substitutions, 14 deletions, and three insertions. These variants were concentrated in two genomic regions involved in plasmid transfer and maintenance (Figure S1), suggesting recombination between plasmids. Another plasmid type, IncFII(pKP91)/IncR, in KpMVS2 harboured *bla*_KPC-2_. All these KPC-encoding plasmids were predicted to be conjugative (relaxase type: MOBF; mating pair formation type: MPF_F), and each carried a variant of the Tn*4401a* transposon harbouring *bla*_KPC-2_ or *bla*_KPC-157_, with 1–2 SNPs between each pair of transposons (Table S1, Figure S1). The comparison between fold-coverages of contigs suggested that each of these three isolates carried a single copy of the KPC-encoding plasmid. Other AMR genes detected in these isolates were chromosomal β-lactamase gene *bla*_SHV_ (variant *bla*_SHV-36_) and fosfomycin resistance gene *fosA10*, which are both intrinsic to *K. pneumoniae* [62–64].

The chromosome of KpMVR1 differed from that of KpMVS1 by six single-nucleotide polymorphisms (SNPs), seven insertions, and eight deletions (including four large deletions illustrated in Figures 1 and S2–5). Seven of these genetic variants were also identified in KpMVS2 (Table 4), which differed from KpMVS1 by 129 SNPs, 13 insertions, and seven deletions. Two large deletions (19.7 kbp and 4.9 kbp) in KpMVR1 could be attributed to an IS-mediated deletion that had previously been observed in *Escherichia coli* (Figures S3 and S5) [65]. Notably, KpMVR1 exhibited deletion of a 54.7-kbp region that comprised operons producing an AcrAB-like multidrug efflux pump and an additional ABC-type Fe^3+^-siderophore transport system in both KpMVS1 and KpMVS2 (Figure 1 and Table S2). Each of the three isolates carried a single copy of the *acrRAB* operon and *tolC* gene, which combine to produce the AcrAB-TolC multidrug efflux pump. However, the permease AcrB in KpMVR1 differed from that in KpMVS1 and KpMVS2 by a destabilising mutation L667R outside the protein’s transmembrane domains.

**Figure 1.**
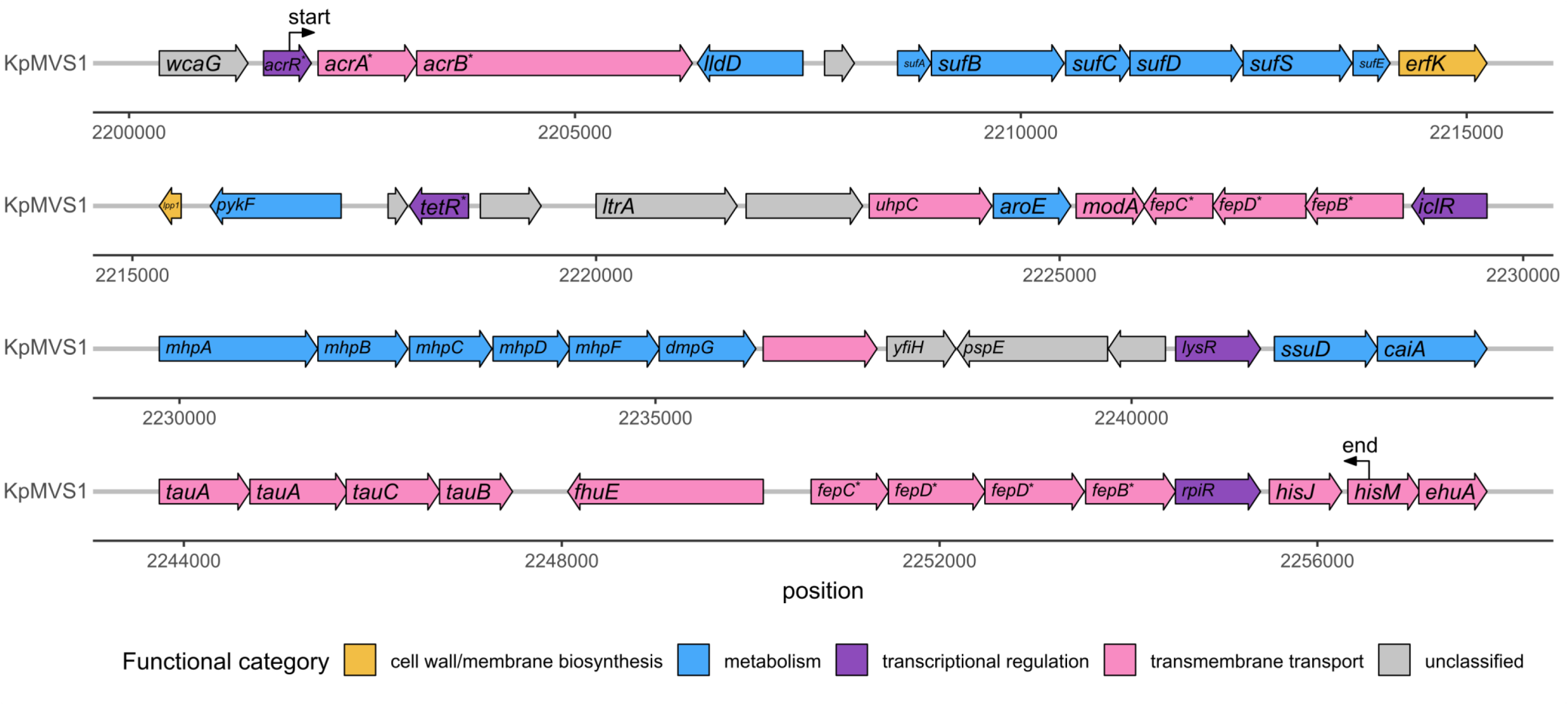
Genetic structure of a 54.7-kbp region in KpMVS1 that was deleted in KpMVR1 (Table 4). Labels “start” and “end” indicate boundaries of the deleted region. Genes without known names are not labelled. Each asterisk indicates an allele from a named gene family.

**Table 4.** Chromosomal genetic variation in isolate KpMVR1 identified via comparison against its progenitor KpMVS1. Coordinates refer to locations in the reference sequence of the KpMVS1 chromosome. Variants shared by both KpMVR1 and KpMVS2 against their common reference sequence of KpMVS1 are indicated by asterisks following the coordinates. The “^” sign indicates an insertion between two consecutive bases in the reference sequence. Abbreviations: CRISPR, clustered regularly interspaced short palindromic repeats; Cas: CRISPR-associated genes; Del, deletion; Ins, insertion; fs, frameshift.

| Location | Locus | Product | Variant type | DNA change | Protein change |
| --- | --- | --- | --- | --- | --- |
| 455179 | *rrl* | 23S rRNA | Substitution | G>T |  |
| 1050491 – 1070213 | Multiple | Type I-E CRISPR-Cas system, *etc.* (Figure S3, Table S3) | Deletion | Deletion of 19,723 bp | Loss of production |
| 1113441 – 1113446 | *flhA* | Formate hydrogenlyase transcriptional activator | Deletion | Deletion of 6 bp | L367Del, T368Del |
| 1218110^1218111 * | *rrl* | 23S rRNA | Insertion | Insertion of base G |  |
| 1578603 | *gyrA* | DNA topoisomerase (ATP-hydrolyzing) subunit A | Substitution | 248C>A | S83Y |
| 1588108^1588109 | *ompK36* | Outer membrane porin OmpK36 | Insertion | Insertion of ISEc68 | Amino acid substitutions |
| 1724677^1724678 * | *xylB* | Xylulose kinase | Insertion | Insertion of base G | I236fs |
| 1792900 | *rfbD* | UDP-galactopyranose mutase | Substitution | 578T>A | M193K |
| 1819375^1819376 | Intergenic |  | Insertion | Insertion of base C |  |
| 2183835^2183836 * | Intergenic |  | Insertion | Insertion of base T |  |
| 2183837 * | Intergenic |  | Substitution | A>T |  |
| 2201799–2256547 | Multiple | Multiple products including transporters (Figure 1, Table S2) | Deletion | Deletion of 54,749 bp | Loss of production |
| 2759671–2759685 | *marR* | Multiple-AMR (Mar) transcriptional repressor MarR | Deletion | Deletion of bases 263–277 | P88–D92Del, K93Q |
| 3157221^3157222 * | Intergenic |  | Insertion | Insertion of base C |  |
| 3189754 – 3223285 * | Multiple | Multiple products (Figure S4, Table S4) | Deletion | Deletion of 33,532 bp | Loss of production |
| 3274833 | *phoQ* | Two-component system sensor histidine kinase | Substitution | C>T | T156I |
| 3580334 – 3585227 * | Multiple | IS3H composite transposon (Figure S5, Table S5) | Deletion | Deletion of 4,894 bp | Loss of production |
| 4104884 | *acrB* | Multidrug efflux RND transporter permease subunit AcrB | Substitution | T>G | L667R |
| 4262704^ 4262705 | *ecpR* | Regulator protein EcpR | Insertion | Insertion (2 bp) | I115fs |
| 4723928 | *rrl* | 23S ribosomal RNA | Deletion | Deletion of base C |  |
| 5349457 | *rrl* | 23S ribosomal RNA | Deletion | Deletion of base C |  |

As for the biosynthesis of siderophores and transport of the iron-siderophore complex, which facilitate cefiderocol to penetrate the OM [66], KpMVS1, KpMVR1, and KpMVS2 were predicted to possess complete enterobactin production and iron-enterobactin transport systems, while none of the yersiniabactin, colibactin, aerobactin, or salmochelin loci were detected, corresponding to a Kleborate virulence score of zero. All three isolates shared the same 19-kbp chromosomal region harbouring a cluster of enterobactin-synthesising genes *entA–F and entH*, enterobactin-exporter gene *entS*, and iron-enterobactin transporter genes *fepA–D* and *fepG*.

Compared with the ciprofloxacin-susceptible isolates KpMVS1 and KpMVS2, KpMVR1 harboured a nucleotide substitution 248C>A in the DNA gyrase gene *gyrA*, resulting in the GyrA mutation S83Y, which is known to reduce ciprofloxacin susceptibility [67]. The three isolates also carried a single copy of the *marRAB* operon. However, KpMVR1 exhibited a unique 15-bp in-frame deletion in the non-essential transcriptional repressor gene *marR* within the *marRAB* operon, causing a loss of five amino acids and an amino acid substitution within the DNA-binding region of MarR (Table 4) [68].

Seven bases at the 5’ end of *ompK36* in KpMVR1 were truncated by an additional copy of insertion sequence IS*Ec68* (three copies in KpMVS1 and KpMVS2, respectively), resulting in a frameshift mutation that replaced the first three amino acids at the N-terminal of OmpK36 with 12 amino acids (Figure 2). The native 21 N-terminal amino acids of OmpK36 encode a Sec-dependent signal sequence (UniProtKB accession: A0A0H3H0Y2) that is required for translocating this protein to the inner membrane of *K. pneumoniae* (Figure 2B) [69]. The signal sequence is subsequently cleaved and OmpK36 is then folded and inserted into the OM (where the protein is functionally active as a porin) in a Bam-complex dependent fashion, a process facilitated by a C-terminal recognition sequence [70]. Whilst *ompK36* from KpMVS1 and KpMVS2 is predicted to encode a complete sec-dependent signal sequence, the 12 amino acids insertion combined with the deletion of three amino acids in OmpK36 from KpMVR1 is predicted to hinder this protein’s translocation to the OM according to the disrupted signal sequence (Figure S6). These predictions were confirmed by polyacrylamide gel electrophoresis of OM preparations, followed by Coomassie staining, that a band corresponding to OmpK36 was present in KpMVS1 but absent in KpMVR1 (Figure 2C). Therefore, the disruption of the Sec-dependent signal sequence of OmpK36 is functionally equivalent to deletion of *ompK36*.

**Figure 2.**
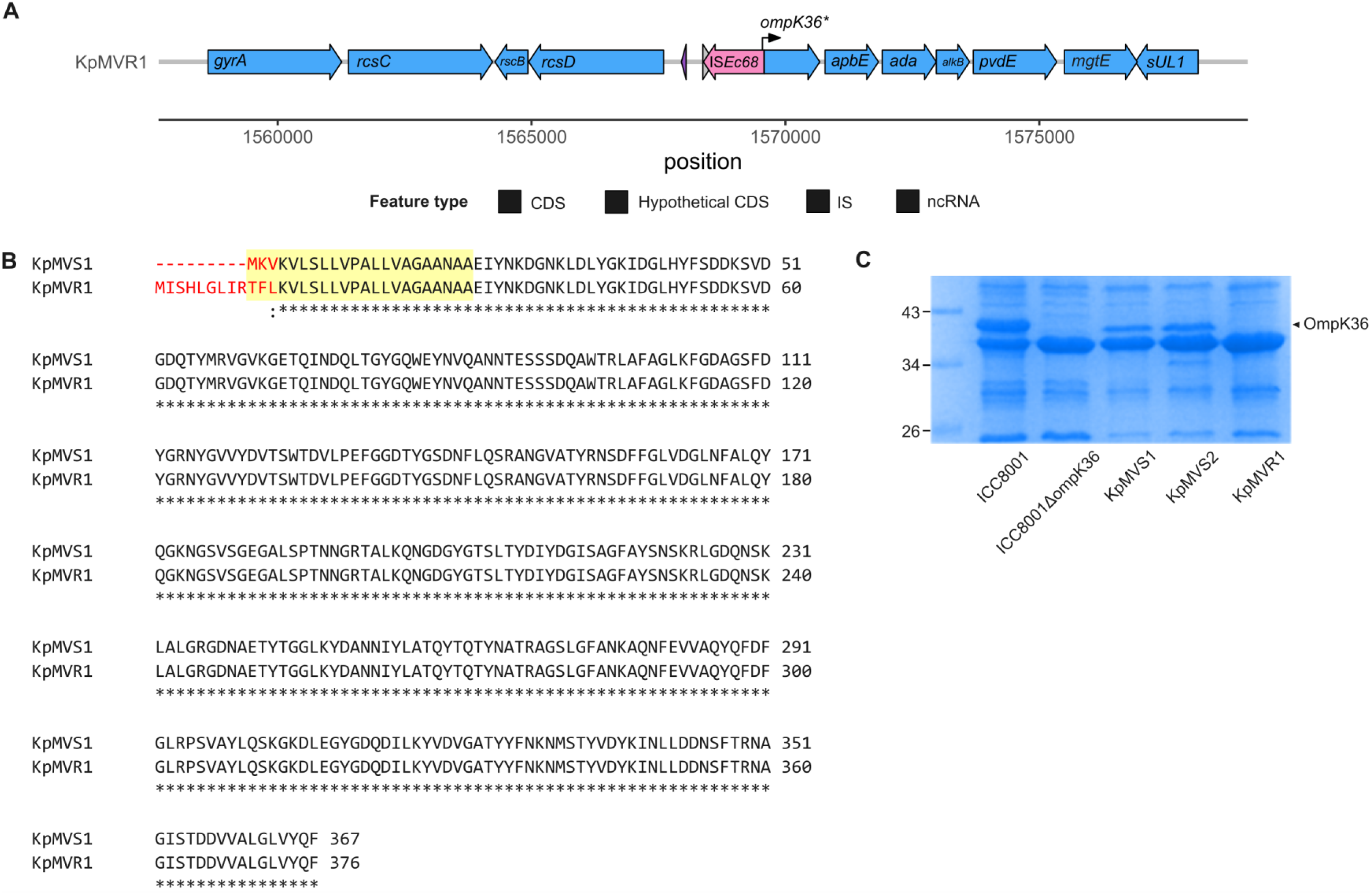
IS*Ec68*-mediated disruption of *ompK36* in isolate KpMVR1. (**A**) Genetic environment of the disrupted *ompK36* in KpMVR1. The arrow labelled “*ompK36\**” denotes the upstream-shifted open reading frame caused by the insertion of IS*Ec68*. Abbreviations: CDS: coding sequence; IS, insertion sequence; ncRNA: non-coding RNA. (**B**) Comparison of predicted OmpK36 sequences using Clustal Omega (www.ebi.ac.uk/jdispatcher/msa/clustalo). Mismatches are highlighted in red, and the 22 N-terminal amino acids signal sequence of OmpK36 are indicated by the yellow shade. (**C**) Coomassie-stained polyacrylamide gel electrophoresis of outer membrane proteins to confirm the absence of OmpK36 in KpMVR1 and the *ompK36*-knockout isolate ICC8001_ΔompK36_.

Regarding the plasmid-encoded TraN proteins, TraN_pKpMVR1_1_ and TraN_pKpMVS2_1_ were identical (NCBI protein accession: WP_049192820.1) and differed from TraN_pKpMVS1_1_ (WP_436914186.1) by six amino acid substitutions (Table S6). Phylogenetic analysis revealed that these proteins belonged to the specialist TraNβ group (Figure S7), which has a narrow host range [50, 71]. Pairwise structural comparison between TraN_pKpMVR1_1_ (TraN_pKpMVS2_1_), TraN_pKpMVS1_1_, and the prototype TraNβ protein TraN_pKpQIL_ showed high consistency (Figures S8), and no amino acid substitution occurred in the characteristic distal β-hairpin (Figure S9), suggesting that the variation in TraN sequences across plasmids pKpMVR1_1, pKpMVS1_1, pKpMVS2_1, and pKpQIL is unlikely to affect the conjugation specificity [72].

### Impact of genetic alterations on antimicrobial resistance

The substitution of *bla*_KPC-2_ with *bla*_KPC-157_ in KpMVS1 (KpMVS1_KPC-157_) and the transconjugant ICC8001_KPC-2_ (ICC8001_KPC-157_) did not affect the susceptibility to meropenem, meropenem-vaborbactam, or imipenem-avibactam but led to a fourfold reduction in the imipenem MIC and a 16- to 32-fold increase in imipenem-relebactam MIC (Table 2). Moreover, this allelic substitution resulted in a 256- to 512-fold reduction in the aztreonam MIC, a 32- to 64-fold reduction in the ceftazidime MIC, and a ≥4-fold reduction in the cefiderocol MIC, but had no effect on the MICs of aztreonam-avibactam or ceftazidime-avibactam. Notably, imipenem and imipenem-relebactam MICs of each KPC-157-producing isolate were identical (Table 2). These findings suggest that KPC-157 has a weaker capacity to hydrolyse imipenem, aztreonam, ceftazidime, and cefiderocol than KPC-2, and that—unlike vaborbactam and avibactam, which inhibit both KPC variants—relebactam inhibits KPC-2 but not KPC-157, which is consistent with a previous report [59].

Knocking out *ompK36* from the ICC8001 chromosome (ICC8001_KPC-2/ΔompK36_ and ICC8001_KPC-157/ΔompK36_) led to a 16-fold increase in the MICs of both meropenem and imipenem, a >33-fold increase in the meropenem-vaborbactam MIC, an 8- to 32-fold increase in the imipenem-relebactam MIC, and a more than twofold increase in the aztreonam MIC (Table 2). These findings are consistent with the role of OmpK36 as an entry route for β-lactams and β-lactamase inhibitors to penetrate the OM [73]. Nevertheless, when comparing MICs of ceftazidime, ceftazidime-avibactam, and cefiderocol before and after knocking out *ompK36* from ICC8001_KPC-2_ and ICC8001_KPC-157_, only two pairs of MICs exhibited notable increases (from 0.125 mg/L to 0.5 mg/L for ceftazidime-avibactam, and from ≤0.06 mg/L to 0.25 mg/L for cefiderocol), while the others showed no appreciable changes, suggesting alternative routes of avibactam’s entry. More generally, the comparison between β-lactam MICs with and without β-lactamase inhibitors for isolates KpMVR1, ICC8001_KPC-2/ΔompK36_, and ICC8001_KPC-157/ΔompK36_ in Table 2 indicates that these inhibitors penetrated the OM via routes other than OmpK36, effectively inhibiting β-lactamases.

The chromosomes of KpMVR1, KpMVS1, and ICC8001 derivatives harboured the same cluster of *ent* and *fep* genes within a 19-kbp region encoding an ABC-type Fe^3+^-siderophore transporter associated with the cefiderocol susceptibility [66]. These isolates did not exhibit any notable difference in cefiderocol MICs despite KpMVR1’s loss of the 54.7-kbp chromosomal region harbouring *fep*-like genes (Table S2), suggesting alternative entry routes of cefiderocol.

## Discussion

In the UK, meropenem-vaborbactam and imipenem-relebactam are recommended for treating adult patients (≥18 years of age) with severe multidrug-resistant infections where therapeutic options are limited, and ceftazidime-avibactam is recommended as an alternative when the disease-causing bacterium produces class D carbapenemase (*e.g.*, OXA-48) [74–76]. Prevalence of resistance in Enterobacterales to any of these three combination antimicrobials was 1–5% across the globe as of 2022 despite regional variation [17, 77–81]. Therefore, the discovery of *K. pneumoniae* isolate KpMVR1, which exhibited unusual resistance to meropenem-vaborbactam, imipenem-relebactam, and ceftazidime-avibactam, in a seriously ill patient is particularly worrisome. The small number of chromosomal SNPs (n=6) and indels (n=10; ≤15 bp each) identified in KpMVR1 when compared with KpMVS1, Patient 1’s exposure to meropenem, meropenem-vaborbactam, and fluoroquinolones, and the unique antibiogram of KpMVR1 in the ICU altogether support the suspected *in vivo* emergence of meropenem-vaborbactam, ceftazidime-avibactam, imipenem-relebactam, and ciprofloxacin resistance in the same *K. pneumoniae* strain during this patient’s hospital stay. A similar shift in the ceftazidime-avibactam susceptibility profile of *K. pneumoniae* has been reported during treatment using meropenem followed by ceftazidime-avibactam [82].

KPC-2 is known to confer carbapenem resistance in Gram-negative bacteria but can be effectively inhibited by vaborbactam, avibactam, and relebactam [83]. Here, we have experimentally determined the effect of carbapenemase KPC-157 on the susceptibility to carbapenems and cephalosporins with or without β-lactamase inhibitors. Our results indicate that KPC-157 does not differ from KPC-2 in its interaction with meropenem or meropenem-vaborbactam. Therefore, the presence of *bla*_KPC-157_ in the single-copy plasmid pKpMVR1_1 alone cannot explain the high-level meropenem-vaborbactam resistance observed in KpMVR1. Notably, KPC-157 appears less capable of hydrolysing imipenem, aztreonam, ceftazidime, and cefiderocol than KPC-2, and is inhibited by vaborbactam and avibactam but not by relebactam.

Isolate KpMVR1 showed genetic changes that may alter the antimicrobial permeability of its OM when compared with isolate KpMVS1. Resulting from an IS-induced sequence disruption, the hindered translocation of OmpK36 to the OM is predicted to hamper the influx of β-lactams and β-lactamase inhibitors into the periplasm, hence elevated MICs of carbapenems and cephalosporins tested in Table 2 with and without β-lactamase inhibitors. Such hampered antimicrobial and inhibitor influx might be further compromised by a decreased expression of *ompK35* in KpMVR1 as a result of the in-frame, presumptively inactivating deletion within the repressor gene *marR* and the consequent upregulation of the *marA* gene [84]. Moreover, the inactivation of *marR* is known to increase the production of the AcrAB-TolC efflux pump, conferring low-level cross-resistance to antimicrobials including β-lactams and ciprofloxacin [85]. However, this upregulation of AcrAB-TolC in KpMVR1 might not alter its antimicrobial susceptibility owing to the possibly destabilised AcrB. Therefore, the high-level ciprofloxacin resistance in KpMVR1 could be primarily driven by the combination of the *gyrA* mutation S83Y and the absence of OmpK36 in this isolate’s OM [67, 86].

This study is limited to three *K. pneumoniae* isolates belonging to the same clone, with only one isolate (KpMVR1) exhibiting elevated MICs of meropenem-vaborbactam, imipenem-relebactam, aztreonam-avibactam, and ceftazidime-avibactam. KpMVR1 harboured multiple AMR-associated genetic alterations. Broader surveillance of genetic variants similar to those identified in KpMVR1 is needed to assess the prevalence and clinical relevance of these putative resistance mechanisms. Although we experimentally validated the individual contributions of *bla*_KPC-157_ and Δ*ompK36* to antimicrobial susceptibility, the genetically reconstructed isolates could not fully replicate the same level of MIC increments as KpMVR1, suggesting that other genetic or regulatory mechanisms may be involved, which remain to be elucidated. Transcriptomic and proteomic profiling could be performed in the future to determine whether regulatory mechanisms also contribute to the observed AMR in KpMVR1.

At the nation level, a review of routine surveillance and reference laboratory samples from 2016-2020 revealed only low levels of resistance to ceftazidime-avibactam in the UK [81]. However, it remains essential that emerging resistance to ceftazidime-avibactam and novel β-lactam/β-lactamase inhibitor combinations is promptly identified and reported through UKHSA’s Second Generation Surveillance System and referral of such isolates to the AMRHAI Reference Unit. Additionally, our study highlights the importance of monitoring the evolving antimicrobial susceptibility profiles of bacterial pathogens within patients during antimicrobial therapy.

## Funding information

This work was mainly funded by the UKHSA. YW is a research fellow funded by the David Price Evans Endowment (grant number: UGG10057) at the University of Liverpool and was an Imperial Institutional Strategic Support Fund Springboard Research Fellow, funded by the Wellcome Trust and Imperial College London (grant number: PSN109). YW, EJ, DM, and KLH are affiliated with the National Institute for Health and Care Research Health Protection Research Unit in Healthcare Associated Infections and Antimicrobial Resistance at Imperial College London in partnership with the UKHSA, in collaboration with, Imperial Healthcare Partners, University of Cambridge and University of Warwick (grant number: NIHR200876). The views expressed in this article are those of the authors and not necessarily those of the NHS, the National Institute for Health Research, or the Department of Health and Social Care.

## Acknowledgments

We acknowledge the Colebrook Laboratory, a facility supported by the NIHR Imperial Biomedical Research Centre (BRC), for providing bioinformatics resources. Part of the bioinformatics analysis was performed on equipment purchased as part of MRC CARP fellowship award MR/T005254/1. We also thank the Institut Pasteur teams for the curation and maintenance of BIGSdb-Pasteur databases at http://bigsdb.pasteur.fr.

## Author contributions

Conceptualisation: KLH and YW; Resources: KLH, JT, GF, FM, NW, and GMR; Methodology: KLH, YW, JLCW, and EJ; Data curation: YW; Investigation and formal analysis: YW, JLCW, JSG, WWL, JT, FM, GMR, KD, IB, GF, EJ, DM, and KLH; Visualisation: YW; Writing – original draft: YW, JLCW, JSG, WWL, GMR, and KLH; Writing – review and editing: all authors.

## Conflicts of interest

Authors declare that there are no conflicts of interest.

## Consent to publish

No sensitive information is disclosed in this manuscript.

## Ethical statement

National surveillance of communicable diseases and outbreak investigation work at UKHSA does not require individual patient consent as per Regulation 3 of The Health Service (Control of Patient Information) Regulations 2002.

## Notes

https://doi.org/10.6084/m9.figshare.29885900

### Competing Interest Statement

The authors have declared no competing interest.

### Summary of Updates

Corrected the following typos in the main text for scientific rigour. 1. Line 71: changed "pathogen transmission" to "resistance transmission". 2. Line 73: changed "administrated" to "administered".

