## Supplementary methods for "Genomic and molecular characterisation of a *Klebsiella pneumoniae* clinical isolate resistant to meropenem-vaborbactam, imipenem-relebactam, and ceftazidime-avibactam"

**August 2025**

### Comparative structural analysis of TraN

Amino acid sequences of $TraN_{pKpMVS1\_1}$, $TraN_{pKpMVR1\_1}$, and $TraN_{pKpMVS2\_1}$ were extracted from the sequence annotations of plasmids pKpMVS1_1 (locus tag: WAS92_RS00545), pKpMVR1_1 (ACNQKT_RS26595), and pKpMVS2_1 (ACNQKS_RS28350), respectively. To contextualise these three proteins, the previously described TraN variants [1] $TraN_{pKpQI}$ (NCBI protein accession: ARQ19727.1), $TraN_{MV2}$ (BAS44060.1), $TraN_{R100\text{-}1}$ (ABD60034.1), $TraN_{pSLT}$ (AAL23498.1), $TraN_{F}$ (WP_000821835.1), $TraN_{MV1}$ (ANZ89826.1), $TraN_{MV3}$ (WP_001398575.1) were downloaded from the NCBI Protein database (www.ncbi.nlm.nih.gov/protein). These 10 amino acid sequences were aligned with the ClustalW algorithm [2], and subsequently, a neighbour-joining phylogenetic tree was generated from the multi-sequence alignment, with the Poisson correction method as implemented in MEGA11 [3]. The phylogenetic tree was visualised using iTOL v7.2 [4].

Three-dimensional structures of $TraN_{pKpMVS1\_1}$, $TraN_{pKpMVR1\_1}$ (identical to $TraN_{pKpMVS2\_1}$), and $TraN_{pKpQI}$ were predicted using AlphaFold 3 [5] with its default parameters on AlphaFold Server (alphafoldserver.com). The top-ranked (model 0) structural models of TraN proteins were visualised using UCSF ChimeraX v1.9 [6]. Superimposition analysis of these models was performed with ChimeraX's Matchmaker tool using default settings, including the use of the "best-aligning" or "bb" chain-pairing method, the Needleman-Wunsch alignment algorithm, and the BLOSUM-62 similarity matrix.
