## Supplementary figures for "Genomic and molecular characterisation of a *Klebsiella pneumoniae* clinical isolate resistant to meropenem-vaborbactam, imipenem-relebactam, and ceftazidime-avibactam"

August 2025

**Figure S1.** Alignment of plasmids pKpMVR1_1 and pKpMVS2_1 against plasmid pKpMVS1_1 using nucleotide BLAST as implemented in Proksee. Plasmids pKpMVR1_1 and pKpMVS1_1 belonged to replicon type IncFII(pKP91)/repB(R1701), and pKpMVS2_1 belonged to IncFII(pKP91)/IncR. The innermost ring represents the number of genetic variants per kbp (calculated using VCFtools v0.1.17) [1] in pKpMVR2_1 compared with pKpMVS1_1. The two middle rings display genes, insertion sequences, and transposons identified in the reference sequence pKpMVS1_1, with arrows indicating orientations of these genetic features (Table S1). Genes without known names are not labelled. The two outer rings show regions of pKpMVR_1 (pink) and pKpMVS2_1 (blue) aligned to pKpMVS1, respectively. This figure was created using Proksee (proksee.ca). "Δ" in gene labels represents a truncated or interrupted feature, and each asterisk represents a variant of an insertion sequence or transposon. Abbreviations: CDS, coding sequence; IS, insertion sequence.

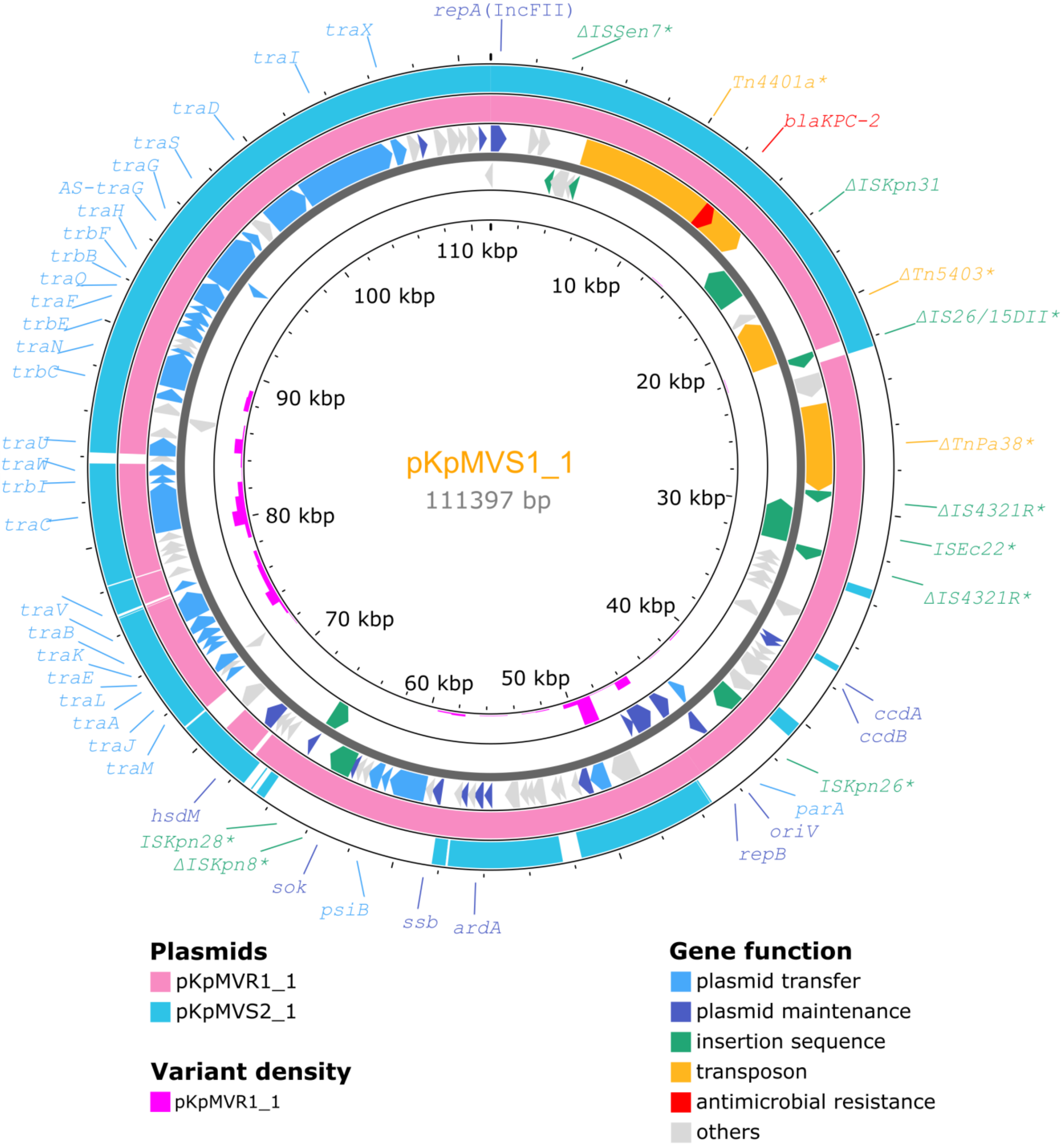

**Figure S2.** A BRIG diagram comparing chromosomes of KpMVR1 and KpMVS2 with that of KpMVS1. Parameters for BLASTn sequence alignment: "-task megablast -ungapped -qcov_hsp_perc 0.8". Four large deletions (>4 kbp) in the chromosome of KpMVR1 when compared to that of KpMVS1 (Table 3) are denoted by digits in filled circles. Genetic structures of these deleted regions are illustrated in Figure 1 and supplementary Figures S3–5.

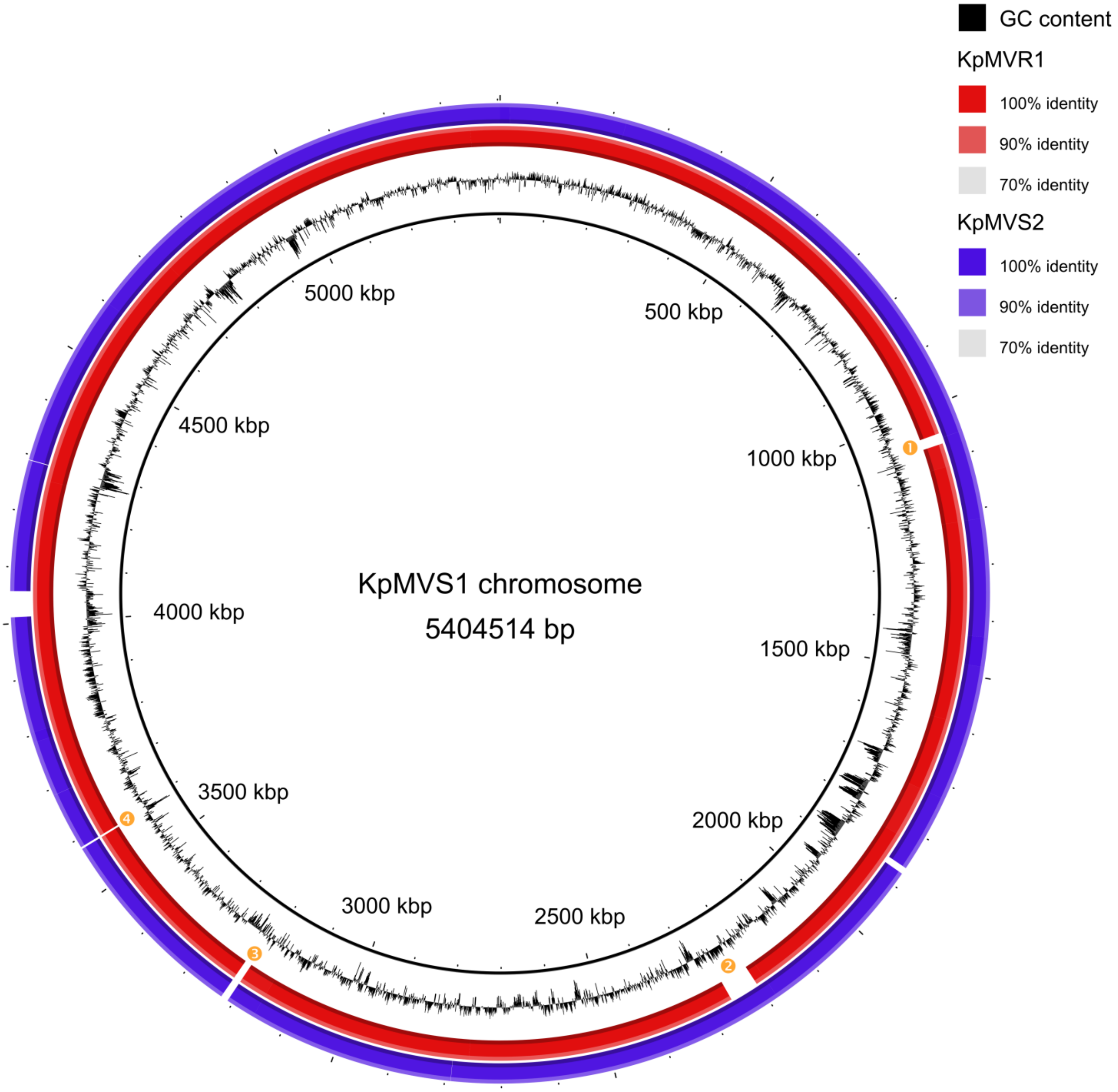

**Figure S3.** Genetic structure of a 19.7-kbp region in KpMVS1 that was deleted in KpMVR1 (Table 4). Labels "start" and "end" indicate boundaries of the deleted region. Genes without known names are not labelled. "IS*1X2**" denotes a variant of the IS*1*-family insertion sequence IS*1X2* (98% nucleotide identity and 100% query coverage). "CRISPR" indicates an array of clustered regularly interspaced short palindromic repeats. See Table S3 for detailed annotations of this region.

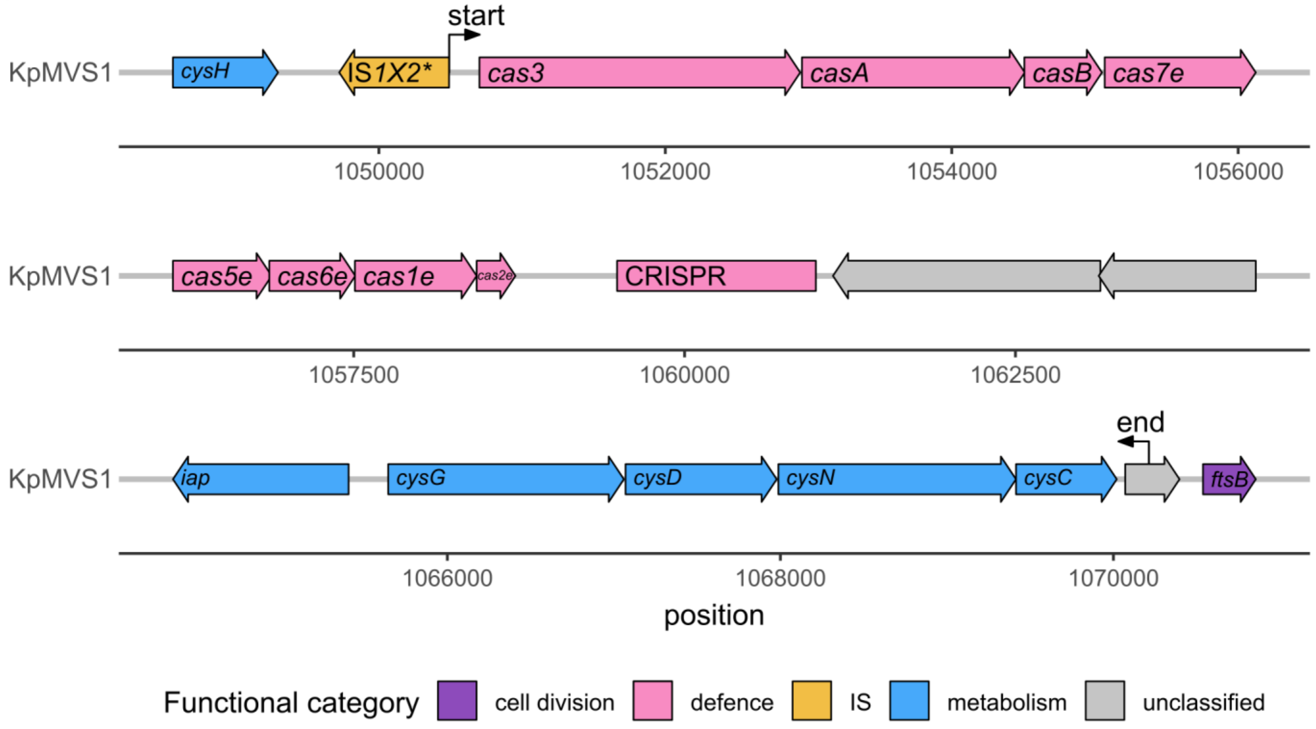

**Figure S4.** Genetic structure of a 33.5-kbp region in KpMVS1 that was deleted in KpMVR1 (Table 4). Labels "start" and "end" indicate boundaries of the deleted region. Genes without known names are not labelled. See Table S4 for detailed annotations of this region.

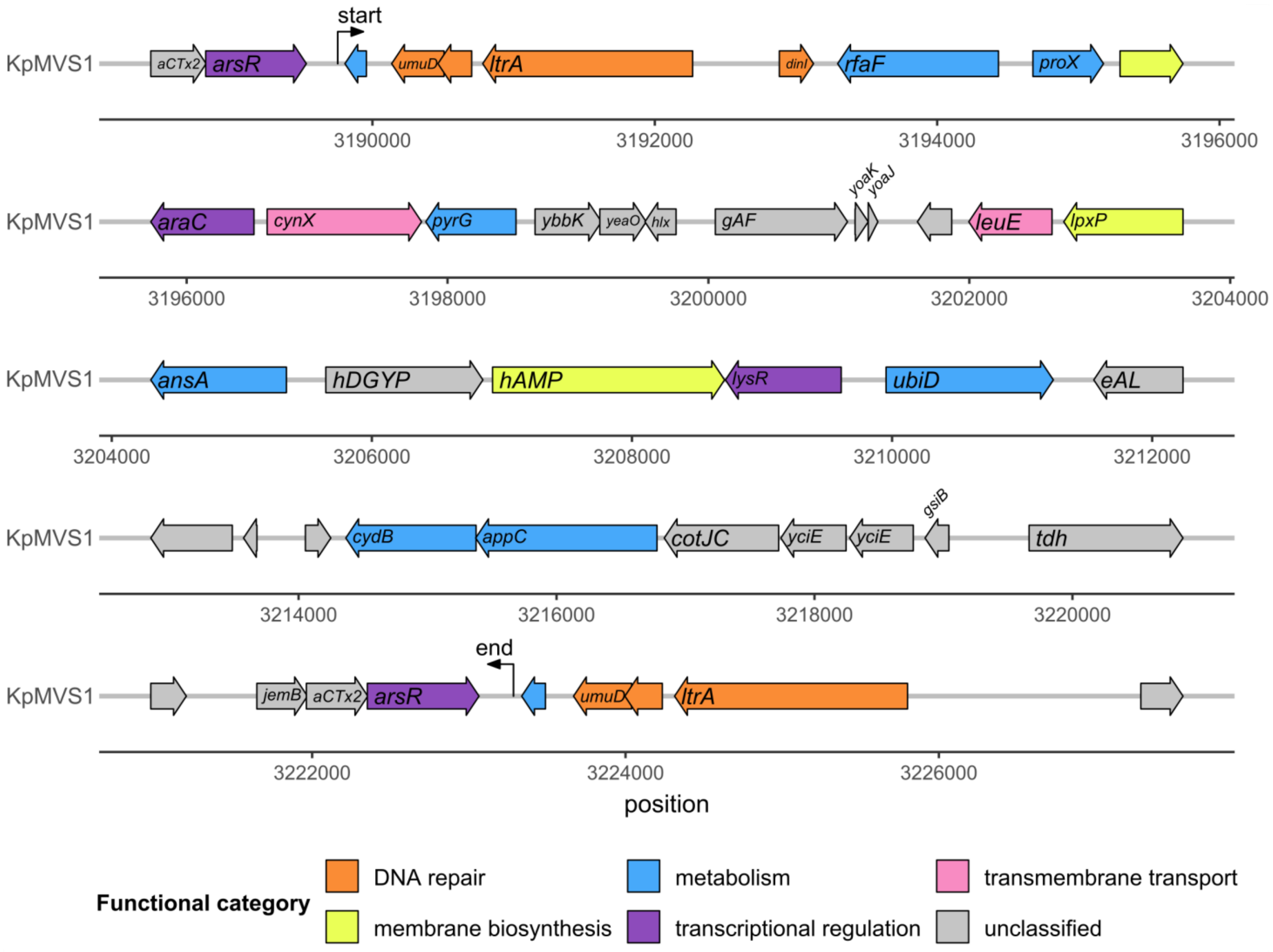

**Figure S5.** Genetic structure of a 4.9-kbp region in KpMVS1 that was deleted in KpMVR1 (Table 4). Labels "start" and "end" indicate boundaries of the deleted region. Genes without known names are not labelled. "IS*3H**" denotes a variant of the IS*3*-family insertion sequence IS*3H* (79% nucleotide identity and 100% query coverage). There were nine copies of this variant in the KpMVS1 chromosome and eight copies in the KpMVR1 chromosome, which is consistent with the content of the 4.9-kbp deletion (Table S5).

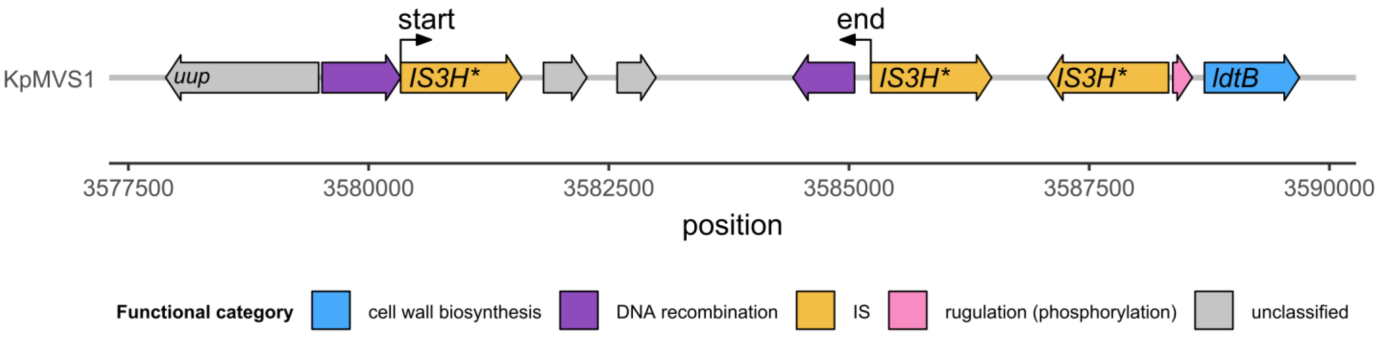

**Figure S6.** Prediction of signal peptides and cleavage site in the wild type OmpK36 (KpMVS1) and its frameshifted variant (KpMVR1), respectively, using the slow model mode of SignalP v6.0. "N", "H", and "C" regions in each protein sequence denote the N-terminal region (Sec/SPI n), centre hydrophobic region (Sec/SPI h), and C-terminal region (Sec/SPI c) of the signal peptide, respectively, whereas "O" denotes the non-signal peptide region (OTHER). The cleavage site (CS) is indicated by the red vertical dashed line. The probability of each amino acid to be part of each peptide region is indicated by a coloured solid curve throughout the protein sequence.

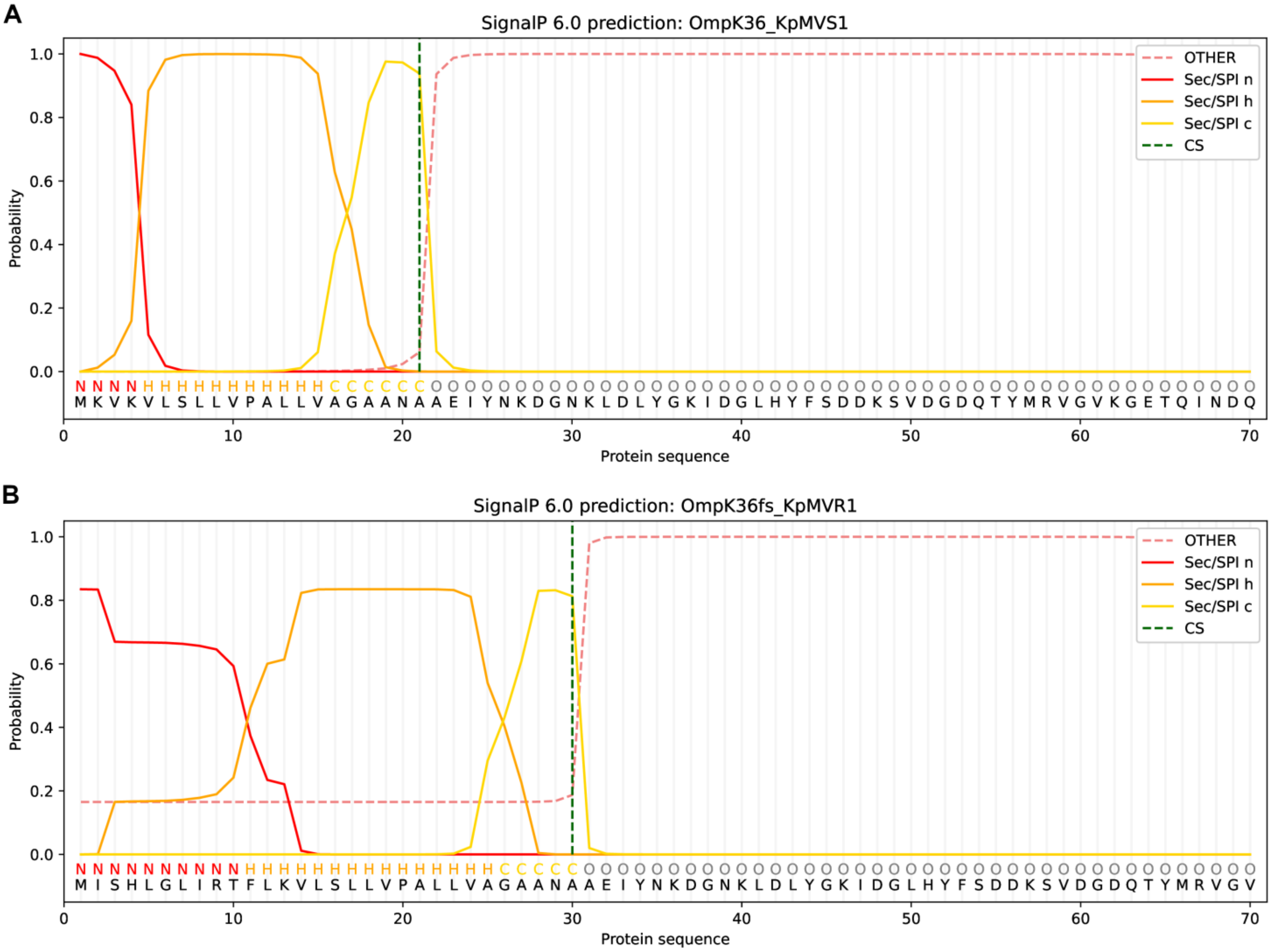

**Figure S7.** Midpoint-rooted neighbour-joining phylogenetic tree of 10 TraN amino acid sequences, where $TraN_{pKpMVR1\_1}$ was identical to $Tra_{pKpMVS2\_1}$. The sequences are named after source plasmids. The structural groups (TraNα, TraNβ, TraNγ, and TraNδ) [2] of TraN are indicated by shaded boxes and labelled. The scale bar represents the number of amino acid substitutions per residue.

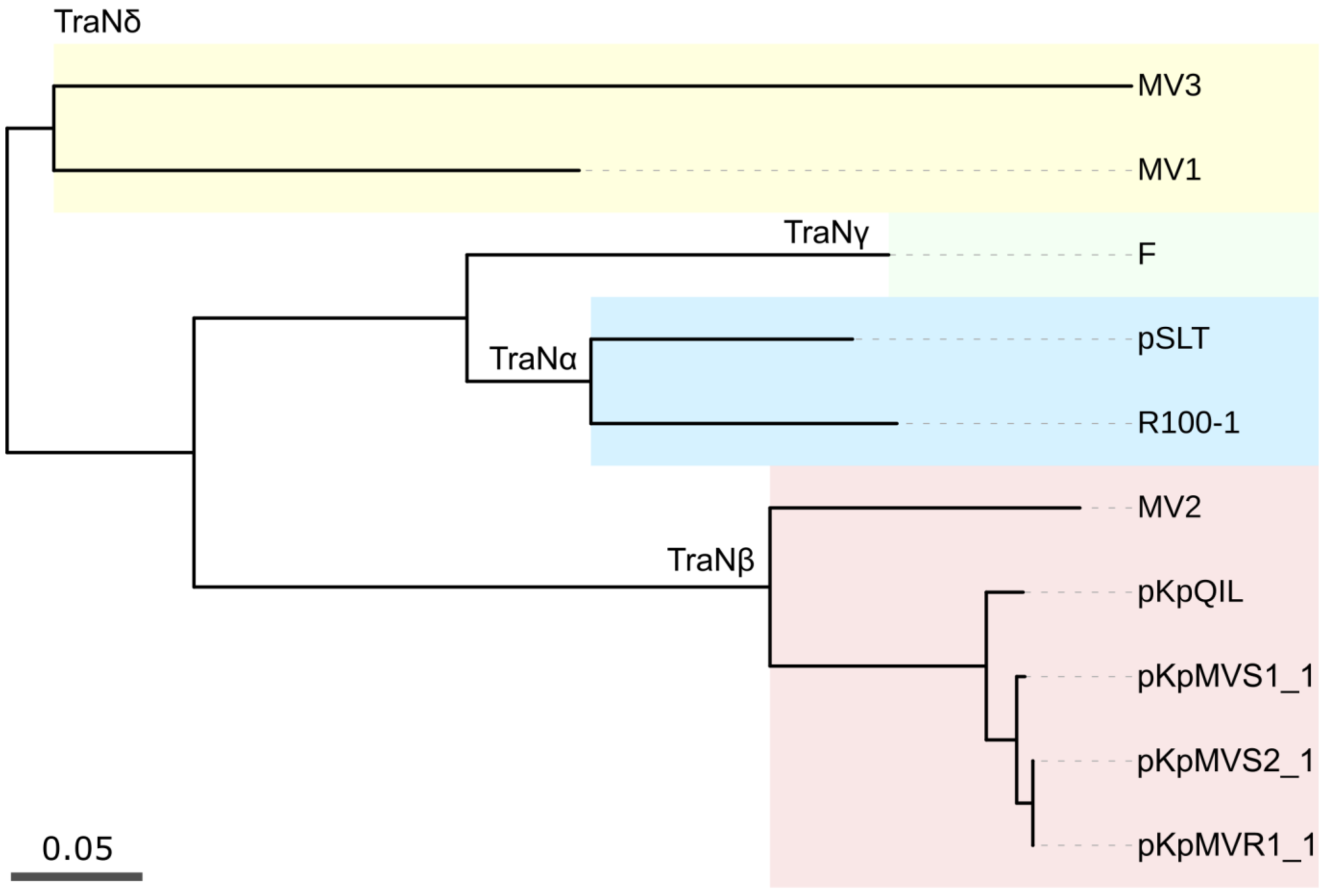

**Figure S8.** Comparison of predicted TraN structures for plasmids pKpQIL, pKpMVS1_1, and pKpMVR1_1. Amino acid chains are represented by ribbons. (**A**) Predicted 3D structures with residues coloured by scores from the predicted local distance difference test (pLDDT). The pLDDT was performed by AlphaFold3 to evaluate the per-residue local confidence of the predicted 3D structure. (**B**) Pairwise structural comparison through the super-imposition analysis. Dashed boxes indicate the tip/sensor domains of TraN [2].

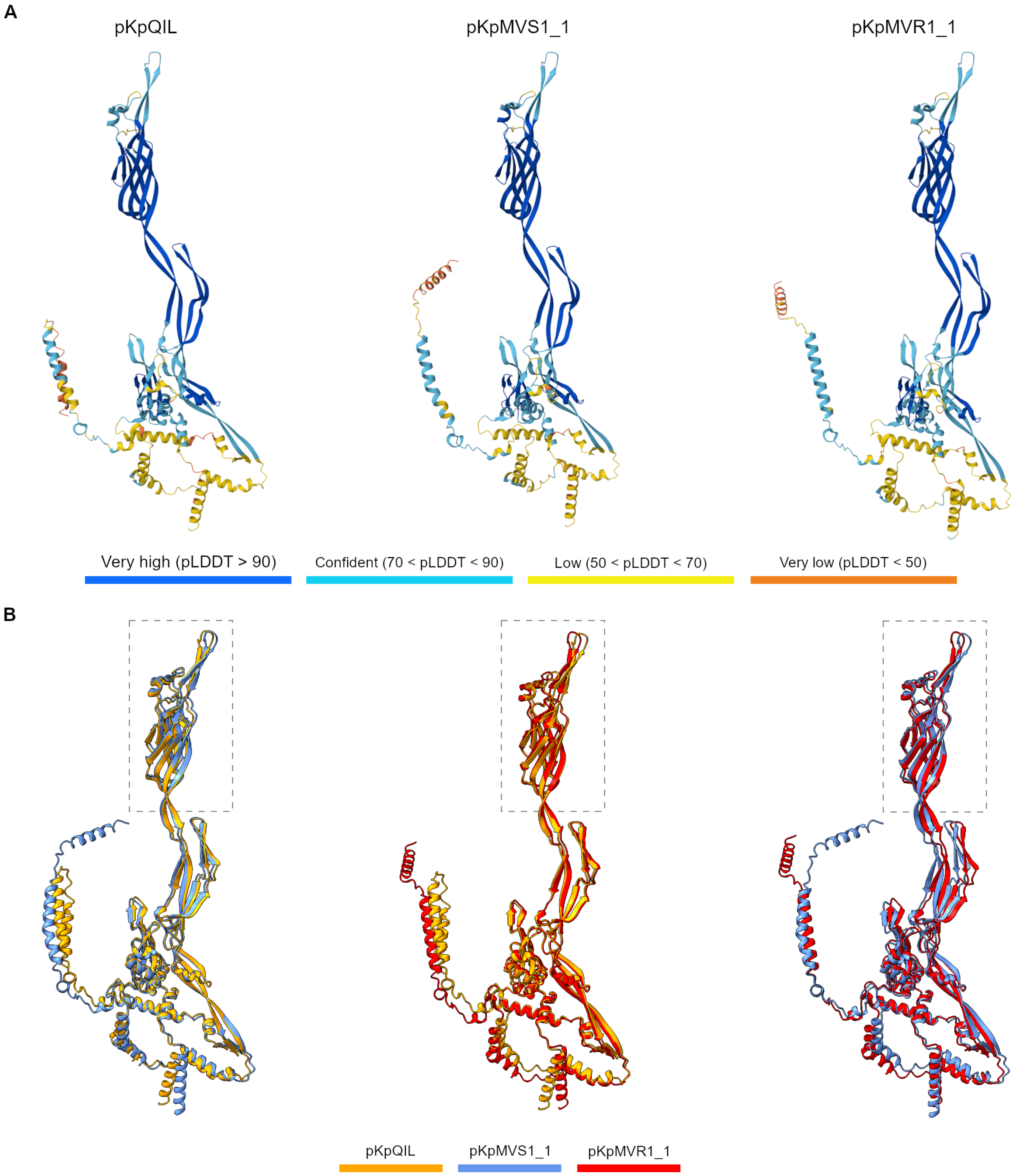

**Figure S9.** Locations of amino acid variation in the tip/sensor domains of TraN from plasmids pKpQIL, pKpMVS1_1, and pKpMVR1_1, with amino acid chains represented by ribbons in the superimposition view. The variable sites are highlighted in yellow, blue, and magenta. In comparison with Figure S8B, the proteins are arbitrarily rotated around the vertical axis to expose all variable sites.

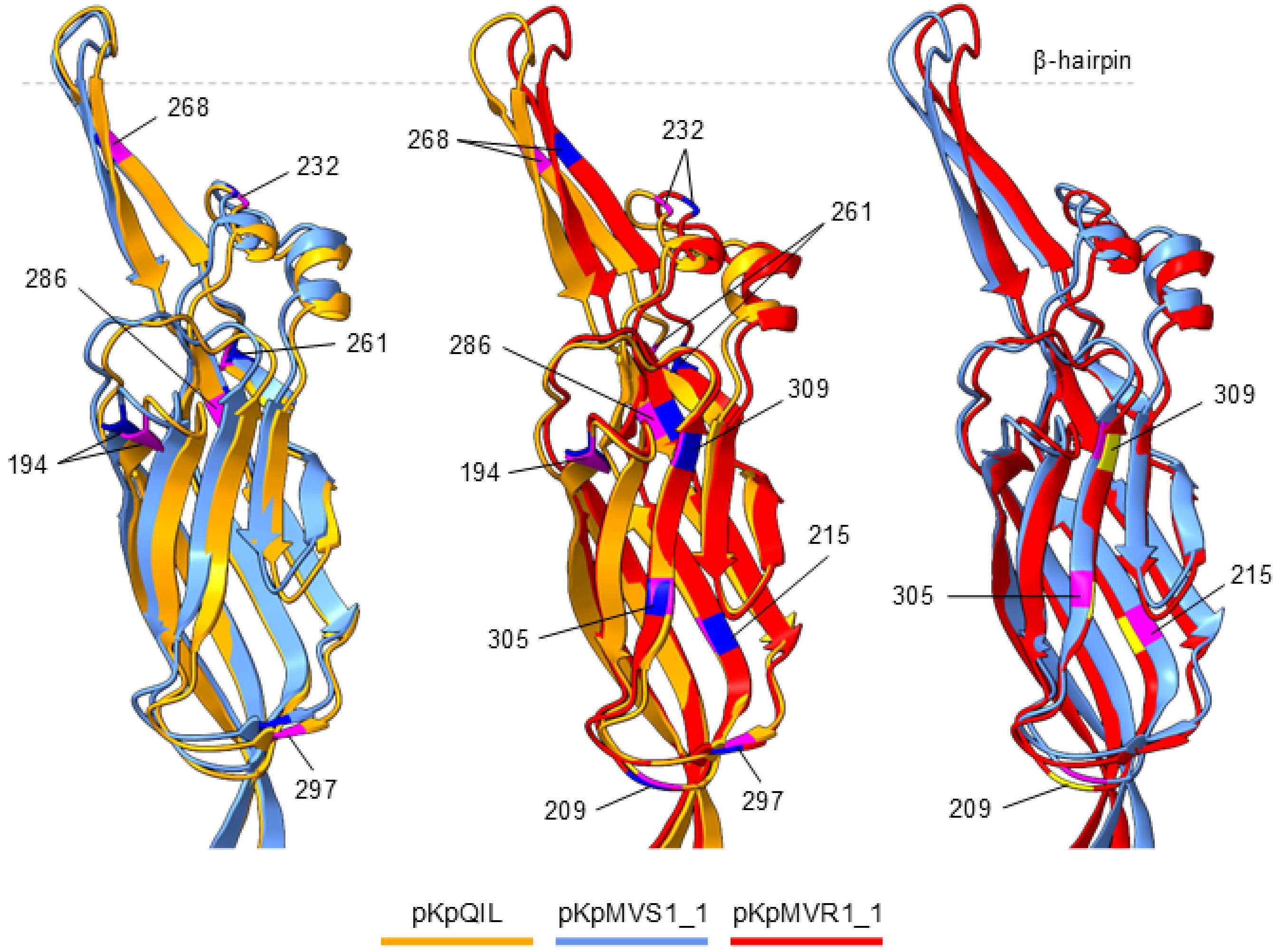
